## Supplementary Information for "Design principles for engineering bacteria to maximise chemical production from batch cultures"

### Supplementary Figures

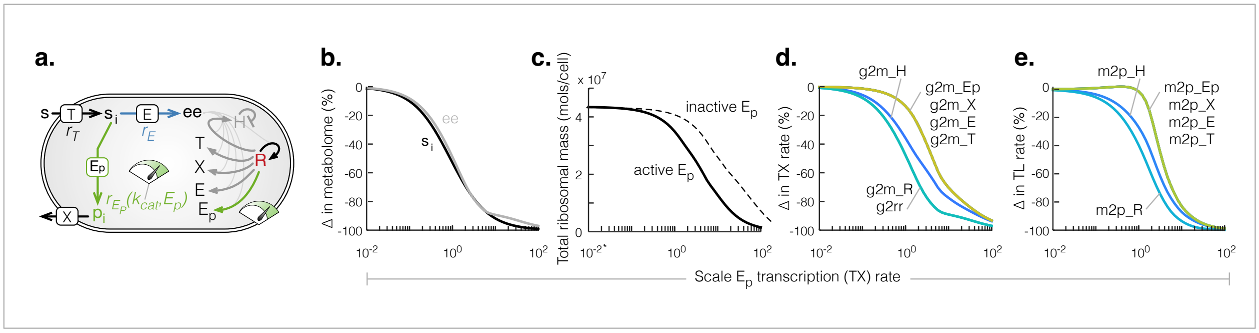


**Supplementary Figure 1: Increasing expression of $E_p$ and drain of $s_i$ increases translation of all enzymes.** (**a**). Schematic of the model indicating our investigation of the effect of increases in the transcription rate (TX) of synthesis enzyme $E_p$. (**b**). % change in the concentration of intermediate metabolite $s_i$ and translation precursor $ee$ for increase in $E_p$ TX rate, compared to zero $E_p$ expression. (**c**). Total ribosomal mass ($R_{active}$ + $R_{inactive}$ + mRNA-R complexes) for increasing $E_p$ TX rate, when $E_p$ is active ($k_{cat} > 0$) or inactive ($k_{cat} = 0$). (**d**). % change in TX rates of enzymes ($T$, $E$, $E_p$, $X$), house keeping proteins ($H$), and ribosomal RNA ($rr$) and ribosomes ($R$), for increasing $E_p$ TX rate. (**e**). % change in translation rate of all mRNA for increasing $E_p$ TX rate.

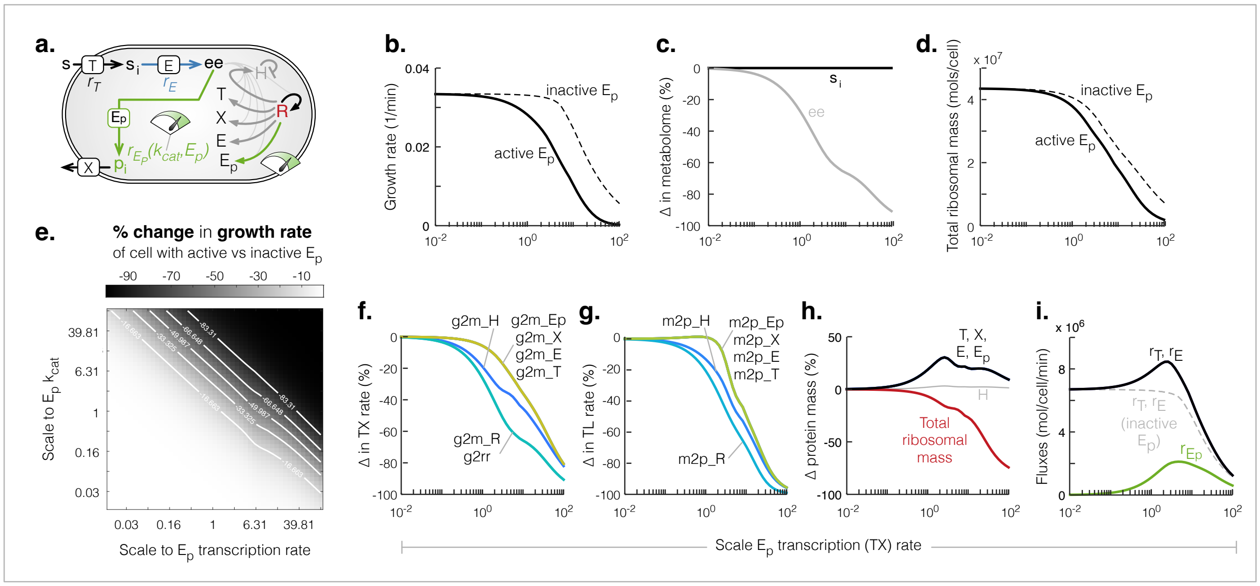


**Supplementary Figure 2: Increasing expression of $E_p$ and drain of translation precursor ($ee$) increases translation of all enzymes.** (**a**). Schematic of the model indicating our investigation of the effect of increasing the transcription rate (TX) and turnover rate ($k_{cat}$) of synthesis enzyme $E_p$ and consequential drain of $ee$. (**b**). Growth rate for increases in the transcription rate (TX) of an active ($k_{cat} > 0$) and inactive ($k_{cat} = 0$) $E_p$. (**c**). % change in the intermediate metabolite $s_i$ and translation precursor $ee$ for increase in $E_p$ TX rate, compared to zero $E_p$ expression. (**d**). Total ribosomal mass ($R_{active}$ + $R_{inactive}$ + mRNA-R complexes) for increasing $E_p$ TX rate, when $E_p$ is active ($k_{cat} > 0$) or inactive ($k_{cat} = 0$). (**e**). Heatmap of the % change in growth rate for increases in the expression of an active $E_p$ ($k_{cat}$¿0) compared to the same level of expression of an inactive $E_p$ ($k_{cat} = 0$). (**f**). % change in TX rates of enzymes ($T$, $E$, $E_p$, $X$), house keeping proteins ($H$), and ribosomal RNA ($rr$) and ribosomes ($R$), for increasing $E_p$ TX rate. (**g**). % change in translation rate of all mRNA for increasing $E_p$ TX rate. (**h**). % change in protein mass for increases in $E_p$ TX rate, compared to zero $E_p$ TX rate. (**i**). Steady state fluxes of all reactions shown in (**a**), for increasing $E_p$ TX rate.

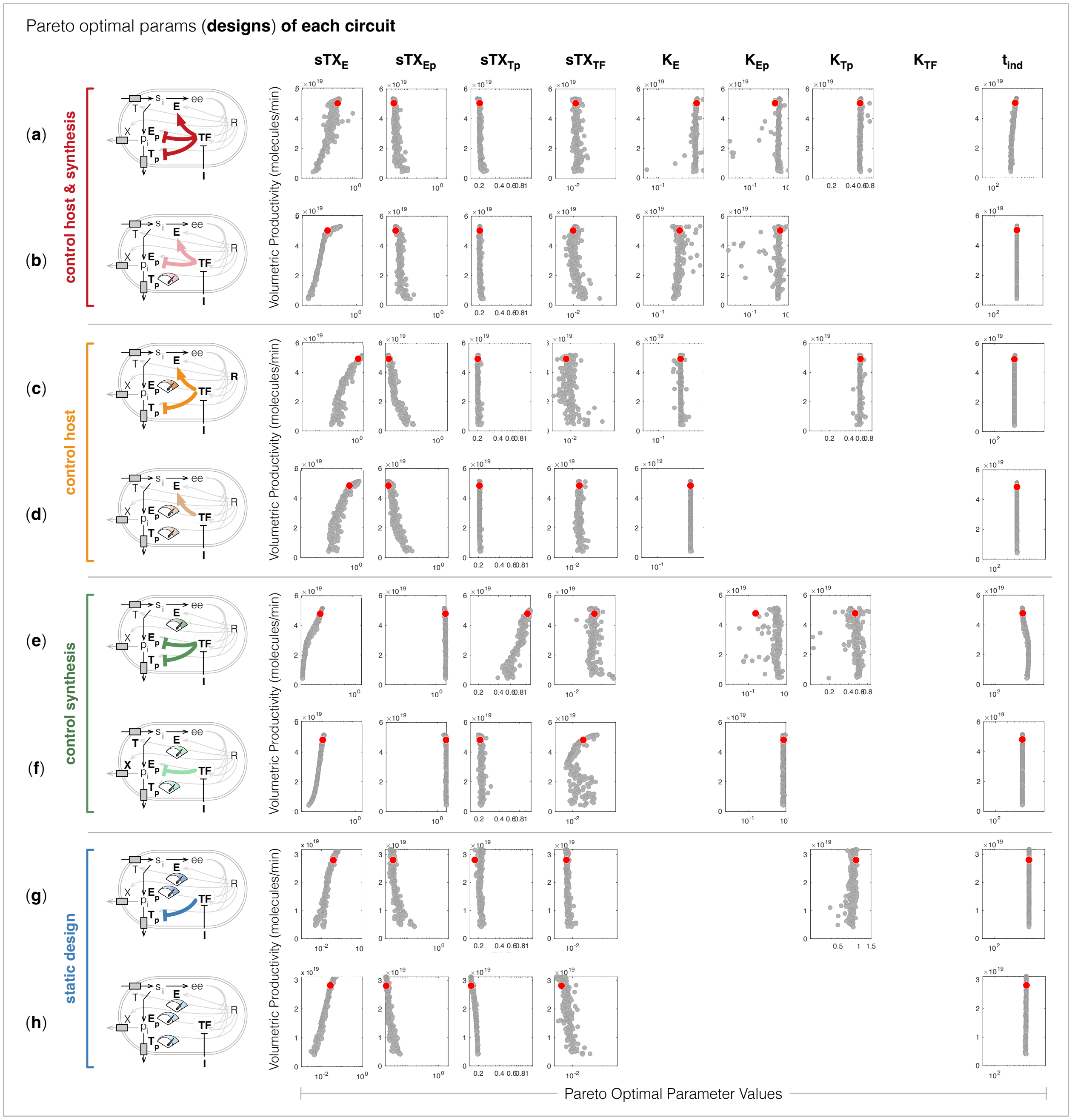


**Supplementary Figure 3: Pareto optimal parameter values, plotted for the respective maximised volumetric productivity,** for circuit topologies representing control on the host and synthesis pathway enzymes (**a** and **b**), control on primarily the host enzyme (**c** and **d**), control on primarily the synthesis pathway enzymes (**e** and **f**), and primarily only tuning the constitutive expression rates of host ($E$) and synthesis pathway enzymes ($E_p$, $T_p$) (**g** and **h**). The parameters (columns) $\mathrm{sTX_E}$, $\mathrm{sTX_{Ep}}$, $\mathrm{sTX_{Tp}}$, $\mathrm{sTX_{TF}}$, represent the coefficients scaling the transcription rates of the enzymes E, $\mathrm{E_p}$, $\mathrm{T_p}$, and TF; $\mathrm{K_E}$, $\mathrm{K_{Ep}}$, $\mathrm{K_{Tp}}$, $\mathrm{K_{TF}}$ represent the "strength" of the TF control on the expression of the respective enzyme, and $\mathrm{t_{ind}}$ represents the time during batch culture when the whole population was induced (by sequestering the TF in each cell).

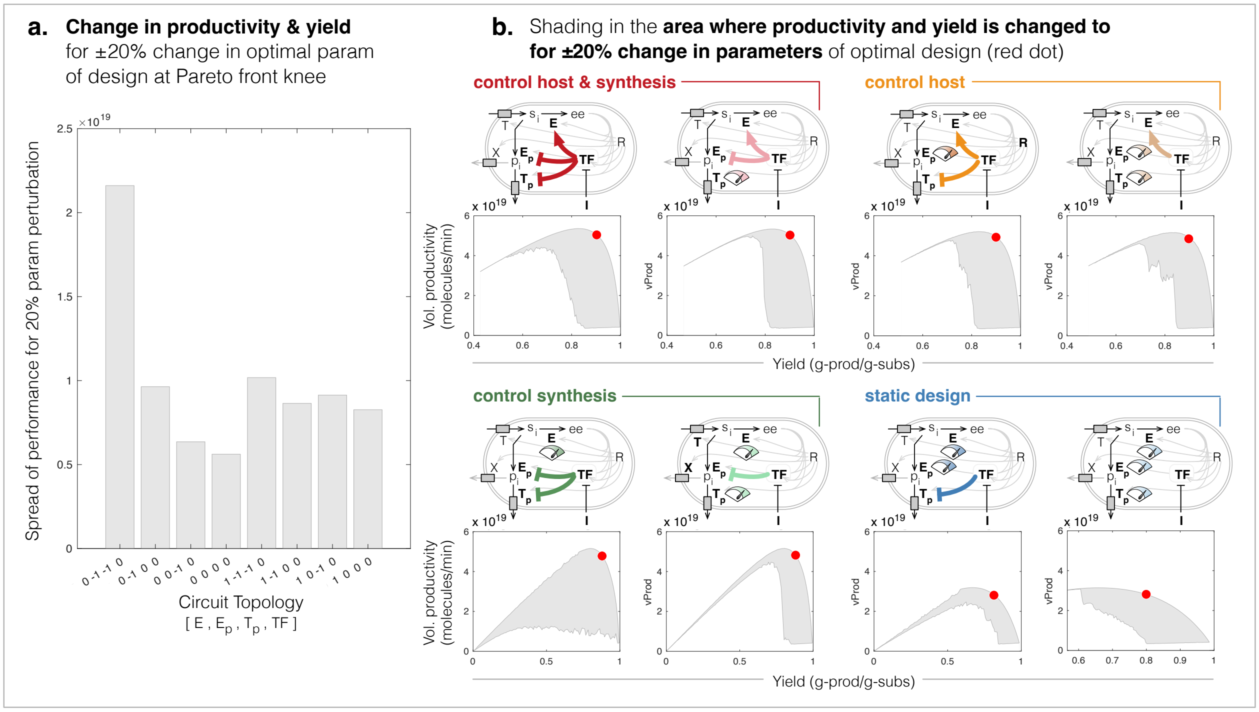


**Supplementary Figure 4: Assessing the robustness of productivity and yield for up to ±20% variation in the Pareto optimal parameters.** (**a**). Bar plot of the measured "area of spread" of the objective values, in the productivity-yield plane, for random variations of up to ±20% in the Pareto optimal parameter values (as defined in Methods 4.4) of the design closest to the optimal ideal point, for the respective circuit topology. (**b**). Plots of the regions (shaded in gray) of the productivity-yield plane to which the two objectives changed to for up to ±20% change in the Pareto optimal parameter value, for each respective circuit topology. The productivity and yield of the original Pareto optimal design is marked by a red dot.

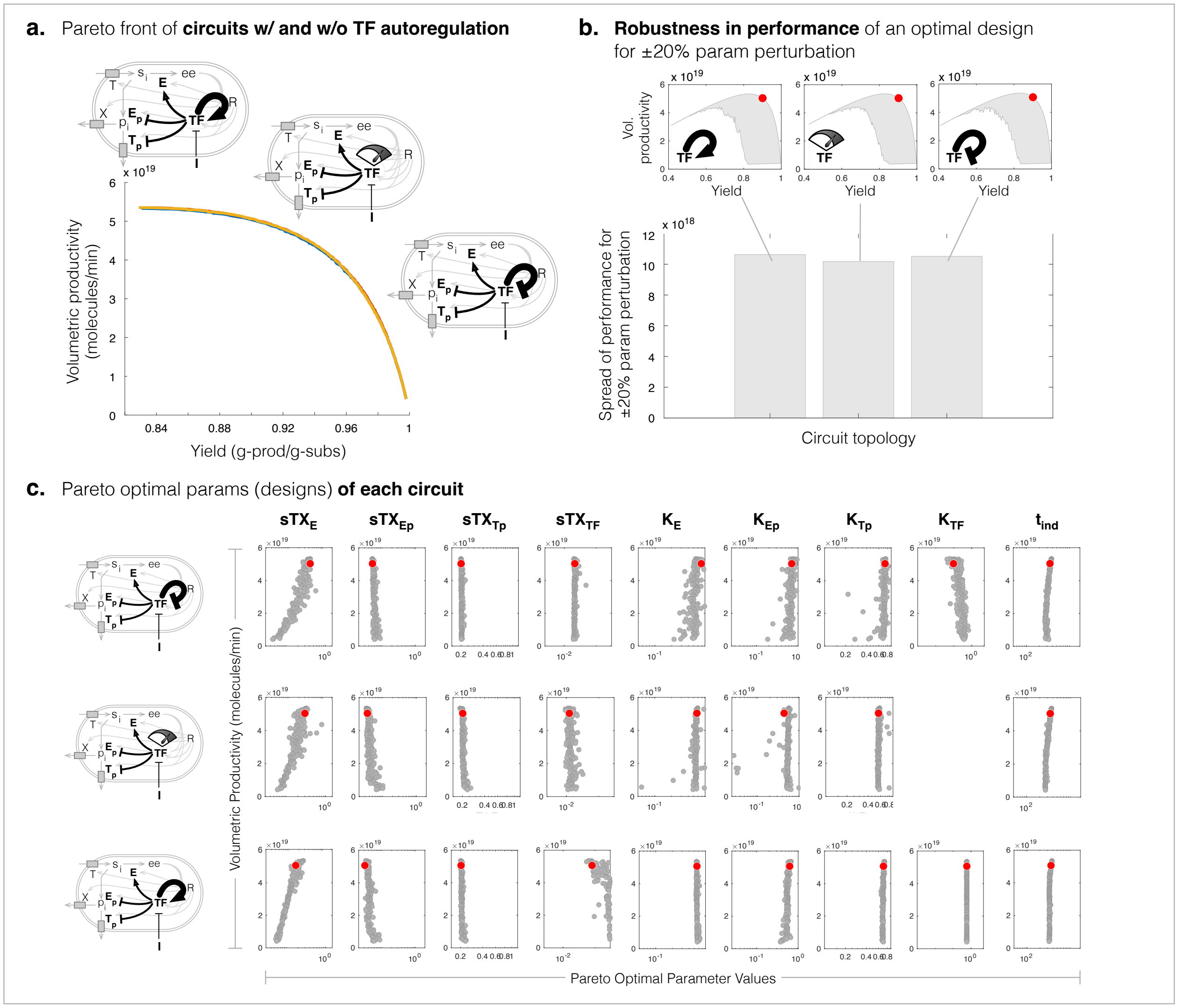


**Supplementary Figure 5: Pareto optimal designs of the dual-control circuit with TF-autoregulation and their robustness to parameter variation.** (**a**). Pareto fronts of the three circuit topologies found on maximising productivity and yield, with the Pareto optimal parameter values shown in (**c**). (**b**). Plots of the measure of "spread" over which the productivity and yield changed to, for up to ±20% variation to the optimal parameter values, of a given optimal design (red dot), of each of the three circuit topologies.

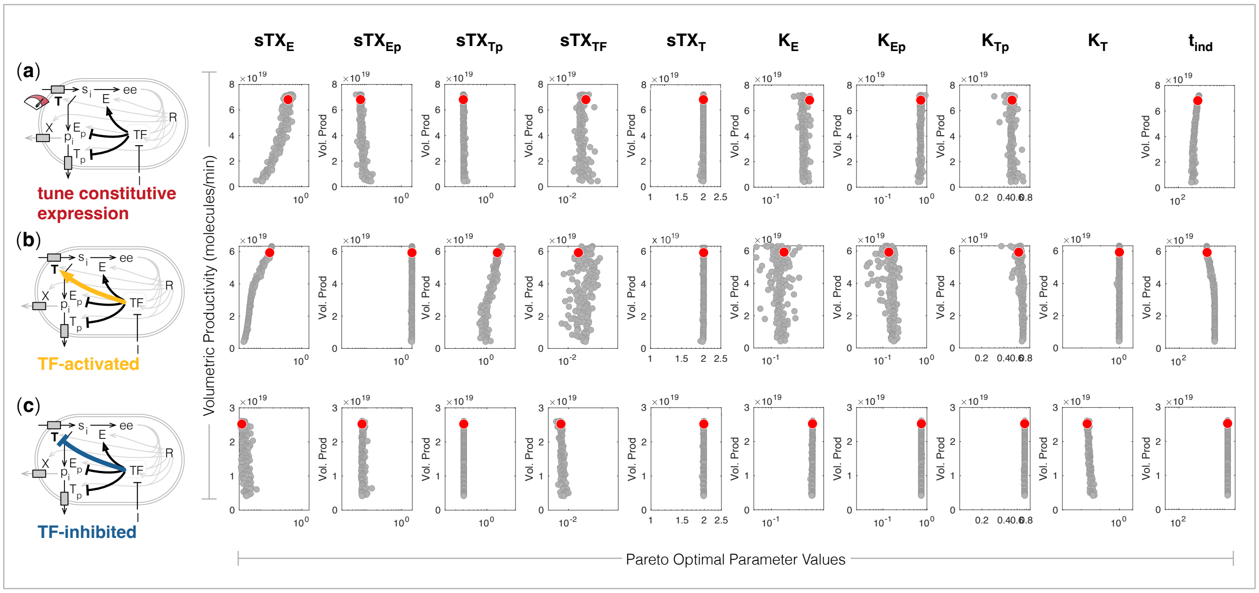


**Supplementary Figure 6: Pareto optimal parameter values of the dual-control circuit augmented with control on expression of nutrient transporter $T$.** (**a**). Pareto optimal designs if constitutive expression of $T$ is tuned. (**b**). Pareto optimal designs if expression of $T$ is deactivated upon inducer addition. (**c**). Pareto optimal designs if expression of $T$ is activated upon inducer addition.

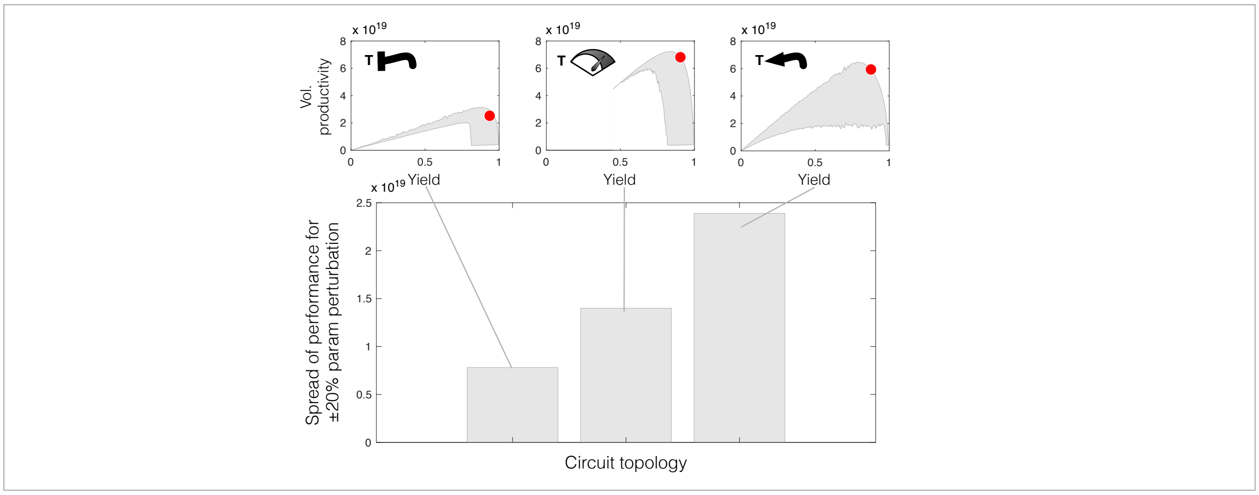


**Supplementary Figure 7: Performance robustness of circuits with control on transporter $T$.** Measuring "spread" of the change in productivity and yield for up to ±20% variation in the optimal parameter values, of the dual-control circuits augmented with three different modes of control on the expression of nutrient transporter $T$: TF-inhibition of T expression (induced activation of T, top left), no regulation but constitutive expression of T (top middle), TF-activation of T (induced deactivation of T, top right).

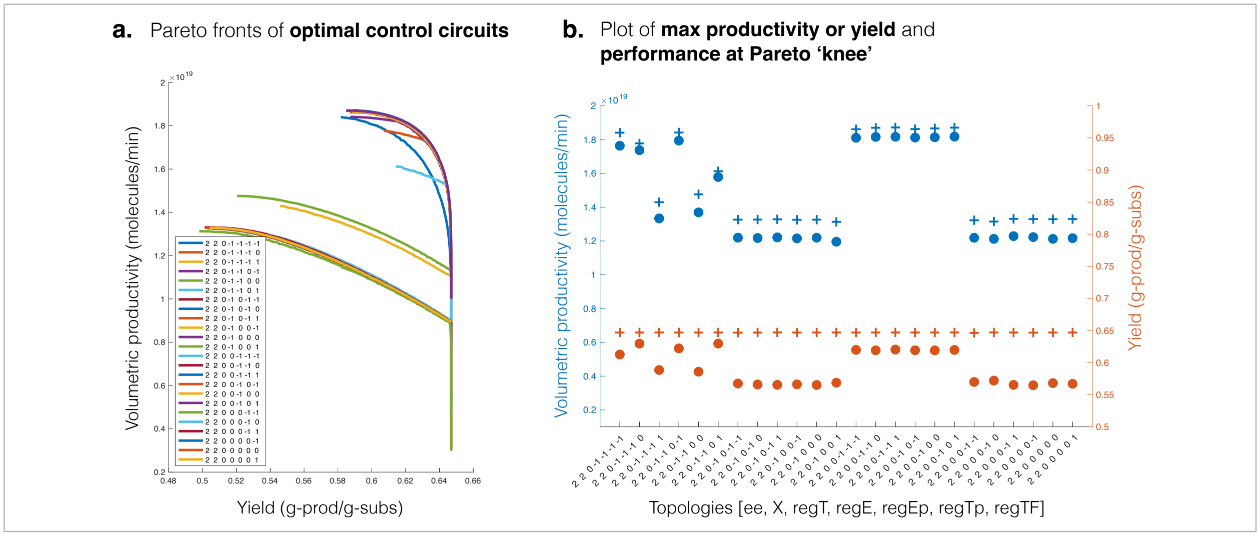


**Supplementary Figure 8: Performance of optimal designs of genetic circuits activating production from translation precursor** "*ee*" (**a**). Pareto fronts of each circuit topology, where the topology is defined by a vector $v$, where $v_1$ indicates product precursor ($2 = ee$), $v_2$ indicates exporters ($2 = X + T_p$), $v_3 - v_7$ indicate uninduced TF-regulation on $T$, $E$, $E_p$, $T_p$, TF (0 = native constitutive expression rate, -1 = inhibition, 1 = activation). (**b**). Plot of the volumetric productivity (blue) and yield (orange) values at the 'knee' of the Pareto front (dots) and their respective maximum values along the Pareto front (plus sign).

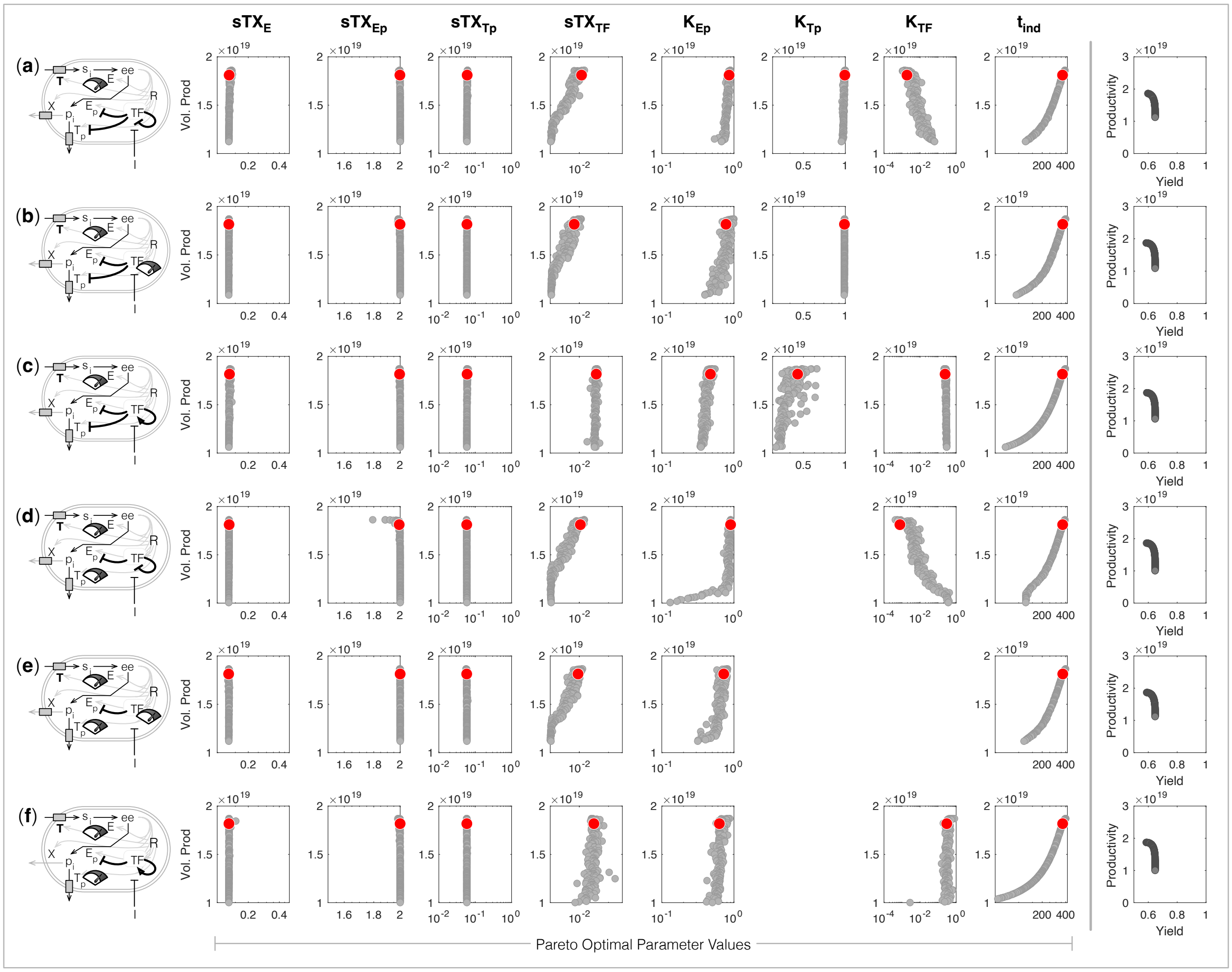


**Supplementary Figure 9: Pareto optimal parameter values of the circuit controlling production from precursor** ***ee*****.** (**a-f**). The six top performing circuit topologies.

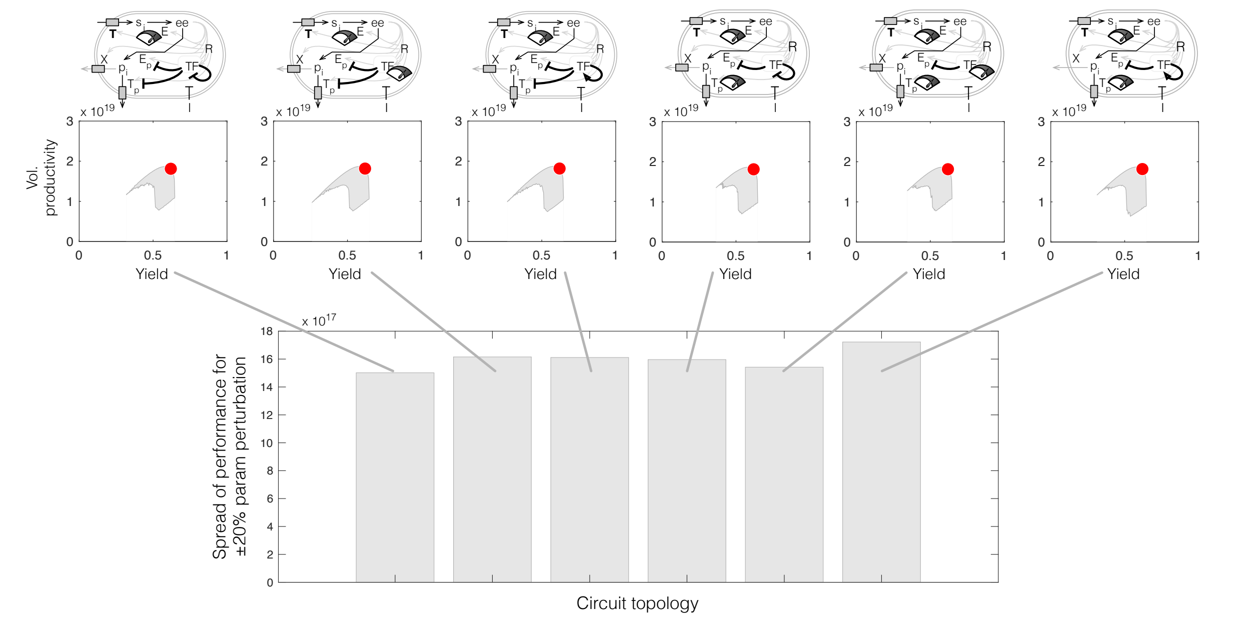


**Supplementary Figure 10: Performance robustness for up to ±20% variation in the Pareto optimal parameters for control on production from *ee*.** Bar plot of the "spread" of the production performance for up to a ± 20% perturbation to circuit parameter values, with the spread illustrated in the top inset plots, for each of the six top performing circuits controlling switch to production from *ee*.

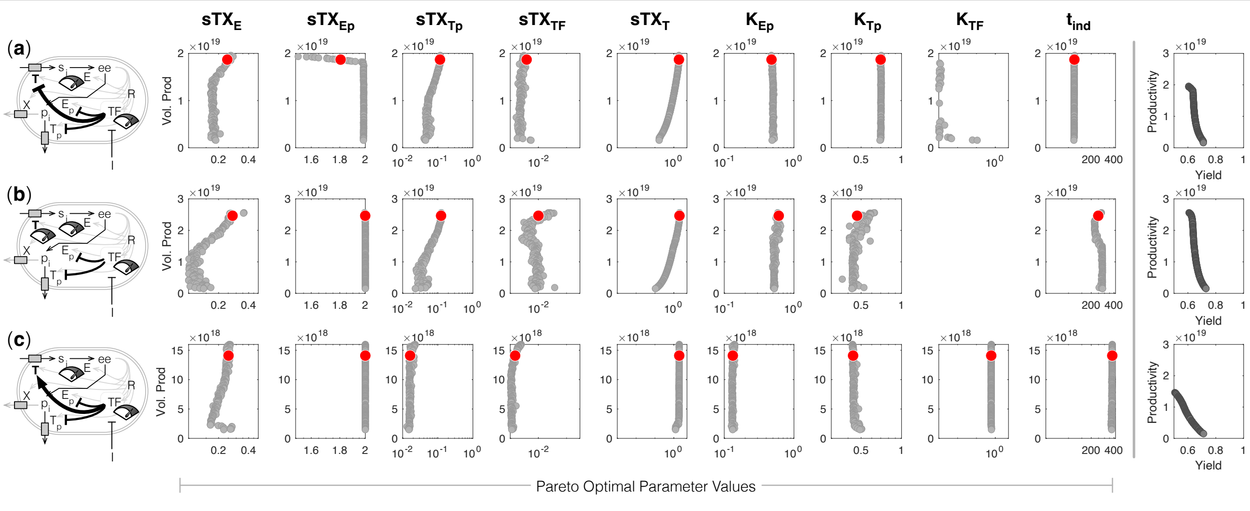


**Supplementary Figure 11: Pareto optimal parameter values of the top performing circuit for switch to production from *ee* augmented with control on expression of nutrient transporter *T*.** Pareto optimal designs (parameter values) if uninduced TF inhibits expression of $T$ (**a**), or has no control on expression of $T$ but $T$'s constitutive expression is tuned (**b**), or if uninduced TF activates expression of $T$ (**c**).

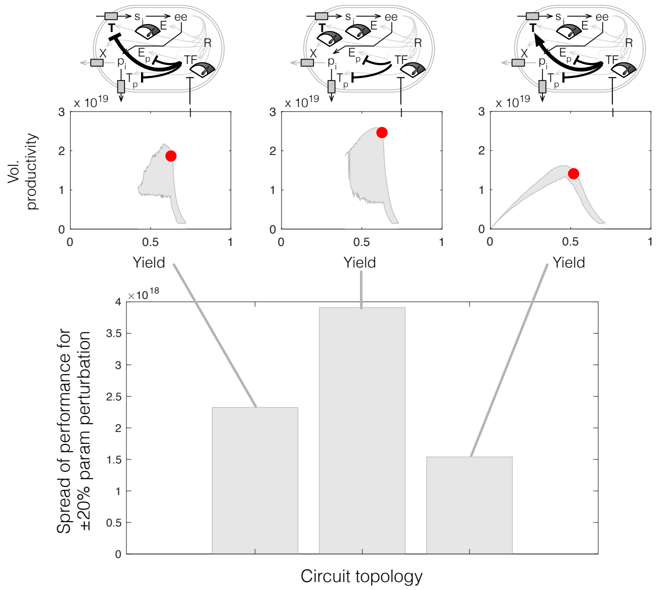


**Supplementary Figure 12: Performance robustness for three circuits controlling production from *ee*.** Bar plot of the "spread" of performance (change in productivity and yield) for up to ±20% change in circuit parameter values, with the "spread" of performance illustrated in the top plots.

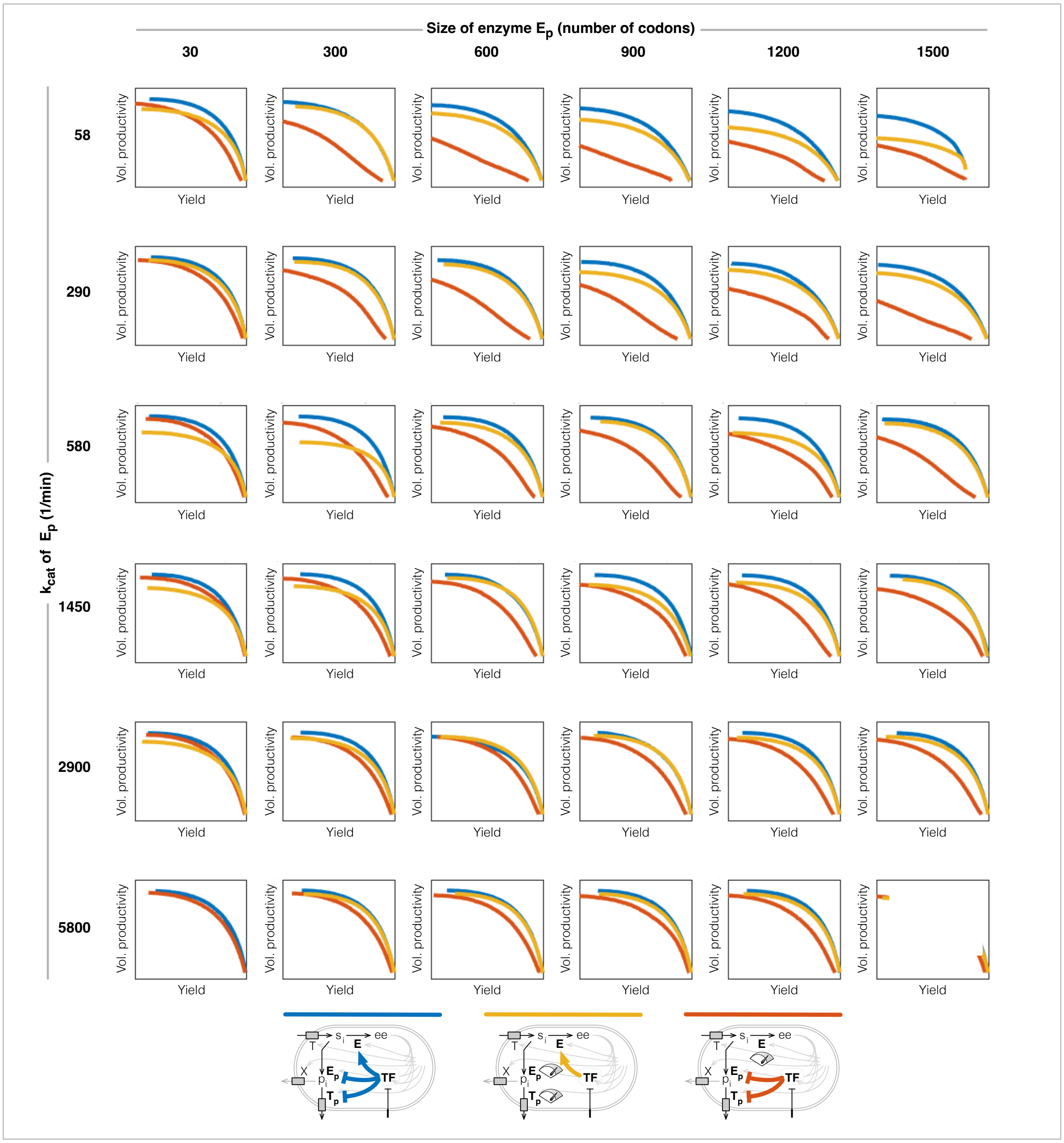


**Supplementary Figure 13: How Pareto fronts change for variation in burden of production pathway enzyme $E_p$.** Pareto fronts found for each of the three core circuit topologies (illustrated at the bottom) controlling production from precursor $s_i$, for variations in the size (number of codons) and turnover rate ($k_{\text{cat}}$) of the product synthesis enzyme $E_p$.

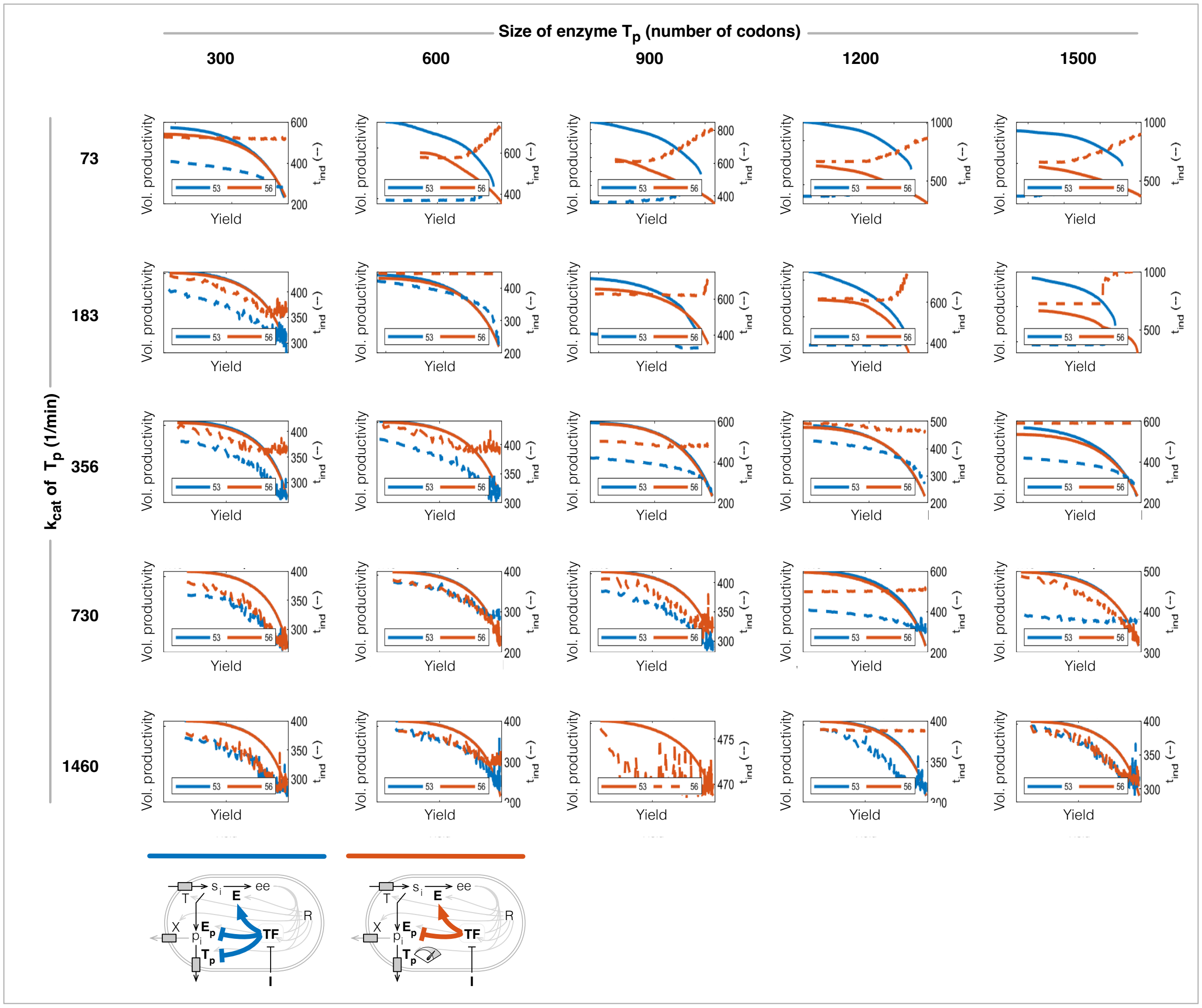


**Supplementary Figure 14: How Pareto fronts change for variation in burden of product transport enzyme $T_p$.** Pareto fronts found for each of the two core circuit topologies (illustrated at the bottom) controlling production from precursor $s_i$, for variations in the size (number of codons) and turnover rate ($k_{\text{cat}}$) of the product transport enzyme $T_p$ (solid curves). The times of induction corresponding to the respective design along the Pareto front (measured by the right y-axis of each plot) is given by the dashed curves.

### SUPPLEMENTARY NOTES

#### SN1 Model description

Bacterial synthesis of non-native chemicals will inherently compete with the cell's native resources. To capture the dynamic interplay between the host and chemical production processes, we use our previously described "host-aware" modelling framework - a coarse-grain computational model of a system of ordinary differential equations describing the dynamics of *E. coli* metabolism, gene expression and growth [1]. Here we describe the host-aware model and how we extended it to account for the expression and competition for host resources of a hypothetical heterologous production pathway, chemical production, the control of enzyme expression with synthetic inducible genetic circuits, and the dynamic environment of batch culture growth. Altogether, this gives a framework of the systems-level, multi-scale view of microbial cell factory performance in batch culture, which captures:

- the competition between the cellular anabolic driver metabolites (e.g. amino acids and ATP) and engineered metabolic pathways,
- the dynamics of these metabolic processes, and their bounding by import/export processes,
- the energy dependence of transcription and translation and the feedback which occurs when the supply of anabolic drivers of these processes is disrupted by an engineered pathway,
- the competition for translational resources occuring between host, pathway and genetic control circuits,
- the impact of metabolic and gene expression burden on host growth rate,
- and the changes in cellular proteome composition (and therefore gene expression rates) which occur when the cell's metabolism and growth are perturbed.

##### SN1.1 The host-aware model of bacterial growth

This non-linear model, also known as a "self-replicator" model, is composed of 16 ordinary differential equations that capture the dynamics of three key cellular processes (illustrated in Supplementary Figure 15):

- **simple metabolism**, describing the uptake of an external substrate to internal $s_i$, and its conversion to an universal energy carrier or translational precursor (e.g., ATP and amino acids), denoted $e$;
- **gene expression** for a coarse-grained genome, consisting of transporters ($T$ denoting host importer, $X$ denoting nonspecific exporters), metabolic enzymes ($E$), and host ($H$) and ribosomal proteins ($R$);

• and **translational resource biogenesis and cell growth**.

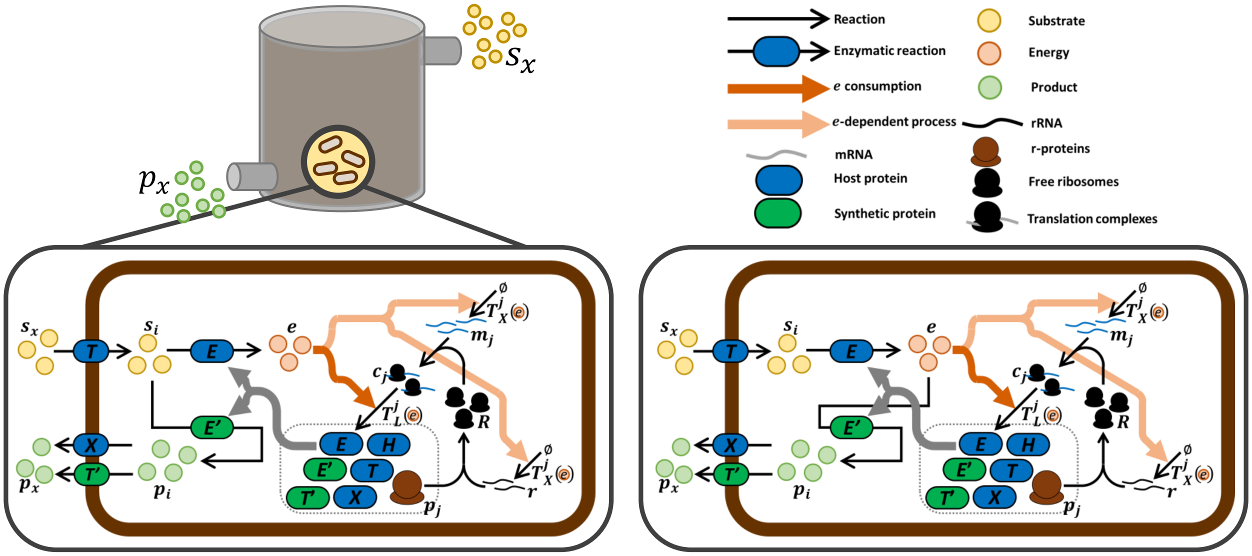


**Supplementary Figure 15: Schematic of the multi-scale model.** The production pathway of chemical $p_i$ is embeded within a host-aware modelling framework. The host-aware framework models a simplified native metabolism composed of internalised substrate $s_i$ and translational precursor $e$, and a proteome composed of host nutrient transporter ($T$), host enzymes ($E$), ribososomes (with r-protein, $xR$, and functional, $R$, counterparts), host "biomass proteins" representing the $q$-fraction from *Scott et al.*, and promiscous host exporter ($X$). The model captures the transcription and translation of the genes of these enzymes/proteins and the utilisation of cellular energy by these processes (denoted $e$). The model is augmented with a heterologous chemical production pathway, catalysed by synthesis protein $E'$ and product exporter $T'$. The host-pathway model is embeded in a larger bioprocess model which captures industrial scale fermentation processes, the production of $p_i$ from the central metabolite $s_i$ (left) or from the anabolic driver $e$ (right).

**Modelling the host's endogenous metabolism.** We consider a simple model of metabolism where an extracellular substrate $s_x$ is imported into the cell to form a pool of internalised substrate $s_i$. This substrate is metabolised to synthesise an translational precursor $e$, yielding $\phi_e$ molecules of $e$ per $s_i$. The species $e$ is then is consumed by translation. All species dilute at a rate $\lambda$ equal to the specific cell growth rate. Applying the law of mass action, the substrate $s_i$ and energy/translational precursor $e$ dynamics within the cell are:

$$\dot{s}_i = \underbrace{\frac{v_T \cdot s_x \cdot T}{K_{mT} + s_x}}_{v_{\text{uptake}}\ \text{Substrate import}} - \underbrace{\frac{v_E \cdot s_i \cdot E}{K_{mE} + s_i}}_{v_{\text{host}}\ \text{Conversion of } s_i \text{ to } e} - \underbrace{\lambda \cdot s_i}_{\text{Dilution}}, \tag{1}$$

$$\dot{e} = \underbrace{\phi_e \cdot \frac{v_E \cdot s_i \cdot E}{K_{mE} + s_i}}_{v_{\text{host}}\ \text{Conversion of } s_i \text{ to } e} - \underbrace{\sum_j \Big(n_j \cdot T_L(c_j,\ e)\Big)}_{\text{Energy consumption by translation}} - \underbrace{\lambda \cdot e}_{\text{Dilution}}. \tag{2}$$

**Modelling gene expression.** We model the expression of nutrient transport and promiscuous export enzymes $T$ and $X$, host metabolic enzyme $E$, host house-keeping proteins $H$, and ribosomal proteins $xR$ (non-active) and $R$ (active). We refer to any of these proteins hereafter as protein $j$. The gene expression model of protein $j$ is composed of an mRNA ($m_j$). This mRNA is born spontaneously at rate $T_{X,j}(e) \cdot F(\cdot)$, where $T_{X,j}(e)$ captures the energy-dependency of transcription of the mRNA of protein $j$ and $F(\cdot)$ is a function capturing any regulation by transcription factors. The mRNA (reversibly) binds to free ribosomes $R$ to produce translation complexes $c_j$. These undergo translation (protein birth) at rate $T_L(c_j,\ e)$ to produce proteins ($p_j$). All species dilute due to cell growth at rate $\lambda$ and mRNAs are also subject to decay at rate $\delta_m$. Applying the law of mass action to this scheme, we derive the following dynamics:

$$\dot{m}_j = \underbrace{T_{X,j}(e) \cdot F(\cdot)}_{\text{Transcription}} - \underbrace{b_j \cdot R \cdot m_j}_{\text{Ribo. binding}} + \underbrace{u_j \cdot c_j}_{\text{Unbinding}} - \underbrace{(\lambda + \delta_m) \cdot m_j}_{\text{Dilution and decay}} , \tag{3}$$

$$\dot{c}_j = \underbrace{b_j \cdot R \cdot m_j}_{\text{Ribo. binding}} - \underbrace{u_j \cdot c_j}_{\text{Unbinding}} - \underbrace{T_L(c_j,\ e)}_{\text{Translation}} - \underbrace{\lambda \cdot c_j}_{\text{Dilution}} , \tag{4}$$

$$\dot{p}_j = \underbrace{T_L(c_j,\ e)}_{\text{Translation}} - \underbrace{\lambda \cdot p_j}_{\text{Dilution}} . \tag{5}$$

**Modelling ribosome availability and its auto-catalysis.** We modelled ribosome synthesis as a two step process; first we considered the transcription of rRNAs ($rr$) and the transcription/translation of r-proteins ($xR$) and secondly, we consider the assembly of the rRNA and r-proteins into the functional ribosome $R$. The dynamics of the rRNA ($rr$) are:

$$\dot{rr} = \underbrace{T_{X,rr}(e) \cdot F(\cdot)}_{\text{Transcription}} - \underbrace{b_\rho \cdot xR \cdot rr}_{\text{Ribo. assembly}} + \underbrace{u_\rho \cdot R}_{\text{Ribo. disassembly}} - \underbrace{(\lambda + \delta_r) \cdot rr}_{\text{Dilution and decay}} . \tag{6}$$

The non-active ribosomal proteins (r-proteins, $xR$) are produced via transcription of their mRNAs ($m_R$) and translation from their translation complexes ($c_R$), their dynamics modelled as:

$$\dot{m}_R = T_{X,R}(e) \cdot F(\cdot) - b_R \cdot R \cdot m_R + u_R \cdot c_R - (\lambda + \delta_{mR}) \cdot m_R, \tag{7}$$

$$\dot{c}_R = b_R \cdot R \cdot m_R - u_R \cdot c_R - T_L(c_R,\ e) - \lambda \cdot c_R, \tag{8}$$

and the dynamics of the expression of ribosomal protein $xR$ modelled as:

$$\dot{xR} = \underbrace{T_L(c_R,\ e)}_{\text{Translation}} - \underbrace{b_\rho \cdot xR \cdot rr}_{\text{Ribo. assembly}} + \underbrace{u_\rho \cdot R}_{\text{Ribo. disassembly}} - \underbrace{\lambda \cdot xR}_{\text{Dilution}} . \tag{9}$$

The dynamics of the functional ribosome are:

$$\dot{R} = \underbrace{b_\rho \cdot xR \cdot rr}_{\text{Ribo. assembly}} - \underbrace{u_\rho \cdot R}_{\text{Ribo. disassembly}} - \underbrace{\lambda \cdot R}_{\text{Dilution}} \ldots \tag{10}$$

$$+\sum_j \underbrace{\underbrace{u_j \cdot c_j}_{\text{Unbinding from mRNAs}} - \underbrace{b_j \cdot R \cdot m_j}_{\text{Binding to mRNAs}} + \underbrace{T_L(c_j,\ e)}_{\text{Translation}}}_{\text{Dynamics of translating proteins } j=\{T,\ E,\ X,\ H,\ R,\ E',\ T', TF\}} \quad ...$$

**Modelling transcription rate.** The birth rate of the mRNA of protein $j$ (transcription) is modelled as an energy-dependent rate with maximal mRNA production rate $\omega_j$ and 'energy-threshold' $o_j$:

$$T_{X,j}(e) = \frac{w_j \cdot e}{o_j + e} \cdot F(\cdot). \tag{11}$$

The $T_{X,j}(e)$ function captures the impact of global regulation; differences in $o_j$ value create differences in response to the cells 'energy status'. Specific gene regulation is captured by $F(\cdot)$ function, otherwise $F(\cdot) = 1$ in the absence of genetic regulation. For host genes $j = \{T,\ E,\ X,\ R,\ rr\}$, we assume no regulation while for generic host 'q-fraction' proteins ($H$), we assume autorepression using a Hill-function:

$$F = 1, \qquad j = \{T,\ E,\ X,\ R,\ r\}, \tag{12}$$

$$F = \frac{1}{1 + (H/k_H)^{h_H}}, \qquad j = H. \tag{13}$$

**Modelling translation rate.** Proteins are born at rate $T_L(c_j,\ e)$ from translation complexes $c_j$ ($j = \{T,\ E,\ X,\ H,\ R,\ E',\ T', TF\}$) at a rate proportional to the cell's internal energy status, $e$:

$$T_L(c_j,\ e) = \frac{\gamma(e)}{n_j} \cdot c_j, \tag{14}$$

where $\gamma(e)$ is a the global peptide elongation rate, i.e. the amino acids incorporated per minute. The $\gamma(e)/n_j$ term yields the protein production rate per translation complex. The global elongation rate is given by:

$$\gamma(e) = \frac{\gamma_{max} \cdot e}{k_\gamma + e}, \tag{15}$$

where $\gamma_{max}$ is the maximal amino acid incorporation rate and $\kappa_\gamma$ is the concentration of $e$ when the peptide elongation rate is half-maximal.

**Modelling growth and burden.** Cellular growth rate is calculated dynamically as a consequence of global protein production rate (as observed in [2] and derived previously in [3]). The instantaneous cellular growth rate ($\lambda$) is:

$$\lambda = (1/M_0) \cdot \gamma(e) \cdot \sum_j (c_j) \tag{16}$$

where $M_0$ is the cellular protome mass, $\gamma(e)$ is the global peptide elongation rate (defined in Eq. 15) and $\sum_j(c_j)$ is the total number of translation complexes. By building the model in this way, we can quantify the effects on host physiology to be assessed by observing the change in this one value.

##### SN1.2 Extending the host-aware model to include chemical synthesis pathway enzymes

We augment the cell model above (Equation (1)-(16)) with equations modelling the dynamics of the one-step intracellular production of a product $p_i$ from either an intermediate host metabolite $s_i$ or expression precursor $e$ (which, below, we denote by general term $m$), catalysed by non-native synthesis enzyme $E'$, at rate $v_{\text{prod}}(m, E')$. The intracellular product $p_i$ is then actively exported from the cell at rate $v_{\text{export}}(\cdot)$. We model the dynamics of the intracellular product formation as:

$$\dot{p}_i = v_{\text{prod}}(m,\ E') - \lambda \cdot p_i - v_{\text{export}}(\cdot). \tag{17}$$

The introduction of the non-native synthesis enzyme $E'$ requires the model to also describe the dynamics of the following associated species: $m_{E'}$, the mRNA of the enzyme; $c_{E'}$, the translation complex of its mRNA and ribosome; and $E'$, the heterologous enzyme itself. The dynamics of this species $\dot{m}_{E'}$, $\dot{c}_{E'}$, and $\dot{E}'$ follow those in Equation (6), Equation (4), and Equation (5), respectively. The extension of the protein set $j$ to include $E'$ updates the dynamics in Equation (2) to account for additional $e$ consumption, updates the dynamics in Equation (10) to account for additional ribosome utilisation, i.e., the competition for host ribosomes, and Equation (16) to account for the impact of translating $E'$ ($c_{E'}$) on growth.

**Modelling the production of $p_i$ from the central metabolite $s_i$.** When considering synthesis of the chemical product from the host intermediate metabolite $s_i$, e.g., a central carbon metabolite, we model the additional drain by $E'$ as:

$$v_{\text{prod}}(s_i,\ E) = (v_{E'} \cdot s_i \cdot E')/(K_{mE'} + s_i), \tag{18}$$

and modify the model of the dynamic availability of $s_i$ (Equation (1)) to:

$$\dot{s}_i = \underbrace{\frac{v_T \cdot s_x \cdot T}{K_{mT} + s_x}}_{v_{\text{uptake}},\ \text{substrate import}} - \underbrace{\frac{v_E \cdot s_i \cdot E}{K_{mE} + s_i}}_{v_{\text{host}},\ \text{conversion of } s_i \text{ to } e} - \underbrace{\lambda \cdot s_i}_{\text{dilution}} - \underbrace{\frac{v_{E'} \cdot s_i \cdot E'}{K_{mE'} + s_i}}_{v_{\text{prod}}}. \tag{19}$$

**Modelling the production of $p_i$ from the translational precursor $e$.** When considering synthesis of the chemical product from species $e$, representing cellular energy or translational precursors, we model its additional drain by $E'$ as:

$$v_{\text{prod}}(e,\ E) = (v_{E'} \cdot e \cdot E')/(K_{mE'} + e), \tag{20}$$

and modify the dynamics of $s_i$ (Equation (2)) to account for $s_i$ conversation into $p_i$ as:

$$\dot{e} = \underbrace{\phi_e \cdot \frac{v_E \cdot s_i \cdot E}{K_{mE} + s_i}}_{v_{\text{host}},\ \text{conversion of } s_i \text{ to } e} - \underbrace{\sum_j \Big(n_j \cdot T_L(c_j,\ e)\Big)}_{\text{energy consumption by translation}} - \underbrace{\lambda \cdot e}_{\text{dilution}} - \underbrace{\frac{v_{E'} \cdot e \cdot E'}{K_{mE'} + e}}_{v_{\text{prod}}}. \tag{21}$$

**Intracellular product export.** We model export of the intracellular substrate (rate denoted $v_{\text{export}}(\cdot)$) in three ways: (i) unlimited export, where the rate of export is limited by the production rate and no specific export protein is required; (ii) the use of a native exporter (a protein denoted $X$) and (iii) the expression and utilisation of a heterologous exporter (protein denoted $T'$). The function $v_{\text{export}}(\cdot)$ takes the following forms:

$$\begin{aligned} v_{\text{export}}(p_i) &= v_{\text{prod}}(\cdot), && \text{unlimited export} \\ v_{\text{export}}(p_i, X) &= (v_X \cdot p_i \cdot X)/(K_{mX} + p_i), && \text{host native exporter} \\ v_{\text{export}}(p_i, T') &= (v_{T'} \cdot p_i \cdot T')/(K_{mT'} + p_i). && \text{heterologous exporter} \end{aligned} \tag{22}$$

Again, when considering the production of the heterologous exporter in addition to the native exporter, the model is extended to also model the dynamics of $m_{T'}$, the mRNA of the heterologous export protein; $c_{T'}$, the translation complex of the exporter mRNA and ribosome; and $T'$, the heterologous exporter protein itself. Their dynamics $\dot{m}_{T'}$, $\dot{c}_{T'}$ and $\dot{T}'$ follow those in Equation (6), Equation (4) and Equation (5), respectively. The extension of the protein set $j$ to include $T'$ updates the dynamics in Equation (2) to account for additional $e$ consumption, updates the dynamics in Equation (10) to account for additional ribosome utilisation and Equation (16) to account for the impact of translating $T'$ on growth.

##### SN1.3 Model of the metabolic controller

We based our control of enzyme expression on a single, chemically-inducible transcription factor, denoted TF. This transcription factor exists in two states: TF, the active DNA-binding form; and $\text{TF}_c$, the non-functional inert complex bound by two molecules of the inducer $I_i$. The inducer is introduced extracellularly as $I$ and we model the rate of its passive diffusion to an internalised form ($I_i$) till concentration equilibrium as:

$$v_{\text{ind}}(I, I_i) = k_{\text{diffI}} \cdot \left(\frac{V_{\text{cell}}}{V_{\text{culture}}} \cdot I - I_i\right). \tag{23}$$

The introduction of protein TF also requires the creation of its associated mRNA and translation complexes with the host ribosomes, $m_{\mathrm{TF}}$ and $c_{\mathrm{TF}}$, respectively, the dynamics of $m_{\mathrm{TF}}$ and $c_{\mathrm{TF}}$ following those in Equation (6) and Equation (4), respectively. Moreover, expressing the additional protein TF means an additional drain on translation precursor $e$ and additional sequestration of the host TF. We thus update Equation (2) and Equation (10) to account for the additional drains. We also update Equation (16) to account for the impact of TF translation ($c_{TF}$) on cell growth. We model the dynamics of the inactive and activated TF (TF and $\mathrm{TF}_c$) availability as:

$$\dot{p}_{TF} = \underbrace{T_L(c_{\mathrm{TF}},\ e)}_{\text{TF protein production}} - \underbrace{\lambda \cdot \mathrm{TF}}_{\text{dilution}} - \underbrace{k_{I,f} \cdot I_i^2 \cdot \mathrm{TF}}_{\text{TF sequestration by } I_i} + \underbrace{k_{I,r} \cdot \mathrm{TF}_c}_{\text{unbinding of } I_i} \tag{24}$$

$$\dot{\mathrm{TF}}_c = \underbrace{k_{I,f} \cdot I_i^2 \cdot \mathrm{TF}}_{\text{TF sequestration by } I_i} - \underbrace{k_{I,r} \cdot \mathrm{TF}_c}_{\text{unbinding of } I_i} - \underbrace{\lambda \cdot \mathrm{TF}_c}_{\text{dilution}} \tag{25}$$

The dynamics of the intracellular inducer are given by:

$$\dot{I}_i = \underbrace{v_{\mathrm{ind}}(I,\ I_i)}_{\text{inducer import}} - \underbrace{2 \cdot k_{I,f} \cdot I_i^2 \cdot \mathrm{TF}}_{I_i \text{ binding to } p_T F} + \underbrace{2 \cdot k_{I,r} \cdot \mathrm{TF}_c}_{\text{unbinding of } I_i} - \underbrace{\lambda \cdot I_i}_{\text{dilution}} . \tag{26}$$

In this work, we considered either the constitutive expression or TF-mediated control (varied strength of inhibition or activation) of the expression of the following key enzymes: the host transporter ($T$), host enzyme ($E$), heterologous pathway enzyme ($E'$), heterologous export protein ($T'$), and the regulating transcription factor itself (TF). To model the impact of TF regulation we modify the regulation function $F(\cdot)$ from Equation (6) for each enzyme in set $j = \{T,\ E,\ E',\ T',\ \mathrm{TF}\}$ to

$$F(\mathrm{TF}) = (1 - \eta^2) + \eta^2 \cdot \left( w_0 + \frac{(K_j \cdot \mathrm{TF})^{\frac{\eta+1}{2}}}{(K_j \cdot \mathrm{TF} + 1)} \right) \tag{27}$$

where $\omega_0$ captures the 'leakiness' of the promotoer, $K_j$ is the transcription factor/DNA affinity rate and $\eta \in \{-1, 0, 1\}$ defines the nature of the TF regulation on the expression of the respective enzyme. We formulated the $F(\cdot)$ function in this way so that when $\eta = 0$ $F(\mathrm{TF}) = 1$ representing constitutive expression; $\eta = 1$ represents a TF enacting activation, before induction, modelled as a Hill function (with cooperativity of 1); and that $\eta = -1$ represents a TF enacting inhibition, before induction, modelled as a Hill function (with cooperativity of 1). Note that as $I$ is an inhibitor of TF binding, $\eta = -1$ results in activation of the target gene upon addition of $I$ (i.e. the repression is removed) and $\eta = +1$ results in a down regulation of the target gene upon addition of $I$ (i.e. the activation is removed).

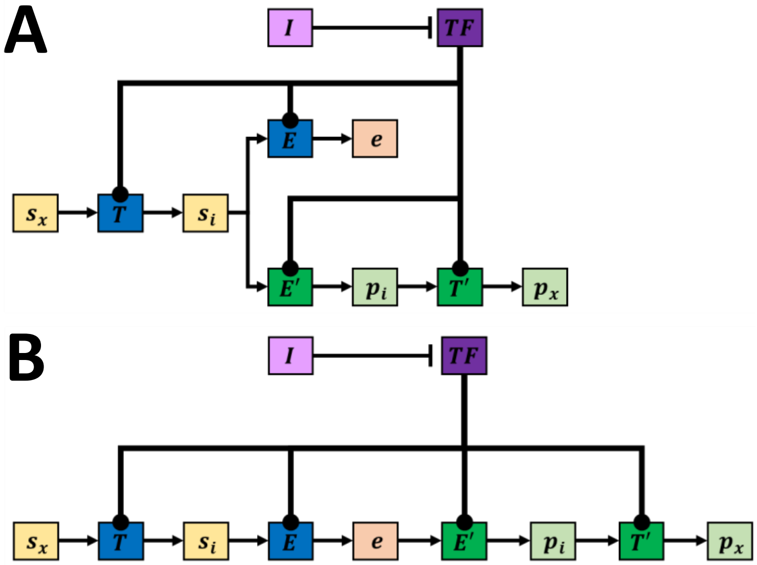


**Supplementary Figure 16: Points of controller action.** The extracellular substrate $S_x$ is imported into cells by the host transporter $T$ to produce intracellular substrate $s_i$. $s_i$ is then converted into the anabolic driver species $e$ (see main text) by the host enzyme $E$. The intracellular product $p_i$ is produced from $s_i$ in panel (A) or from $e$ in panel (B). This reaction is catalysed by the heterologus enzyme $E'$. $p_i$ is exportered from the cells by the heterologous exporter $T'$. Note we do not show the topologies where $p_i$ export is not limited or where a native exporter is used in this figure. The small molecule inducer $I$ acts to inhibit the action of the transcription factor $TF$. The transcription factor can act in an activator or inhibitory on $T$, $E$, $E'$ and/or $T'$. (A) The controller network for a pathway where $p_i$ is produced from $s_i$. (B) The controller network for a pathway where $p_i$ is produced form $e$.

##### SN1.4 Modelling population growth and the external environment

The "cell model" is composed of metabolism, gene expression and growth dynamics, given by Equation (1)-Equation (16) and augmented with pathway and circuit models (Equation (17)-Equation (27)), is embedded within a four state model of batch fermentation. The batch fermentation model consists of the cell population $N$, the extracellular substrates $S$, the extracellular product $P$ and inducer $I$. The dynamics of the extracellular substrate ($S$) take up by the population of cells, at rate $v_{\text{uptake}}$, is modelled as:

$$\dot{S} = -v_{\text{uptake}}(S, p_T) \cdot N. \tag{28}$$

The extracellular product produced from the cells, at rate $v_{\text{export}}$, is modelled as:

$$\dot{P} = v_{\text{export}}(\cdot) \cdot N. \tag{29}$$

The passive diffusion of the inducer is imported into the cell, where the import is balanced against a building intracellular concentration, and modelled as:

$$\dot{I} = v_{\text{ind}}(I,\ I_i) \cdot N. \tag{30}$$

The total population ($N$) grows at rate $\lambda(e,\ c_j)$ (Equation (16)) and is modelled as:

$$\dot{N} = \lambda(e,\, c_j) \cdot N. \tag{31}$$

The population dynamics follow the expected logistic growth curve of batch culture: as the population $N$ grows, substrate $S$ is depleted and so $\lambda(\cdot)$ calculated by the cell model decreases over time, eventually reaching zero when the extracellular and intracellular substrates reach zero.

##### SN1.5 Table of model parameters

**Supplementary Table 1:** Model parameters, table 1 of 2. Column "Range" defines range ([...]) or set ({...}) of values explored when solving optimisation problems defined in Methods 4.3. "mols" = molecules.

| Parameter | Description | Nom. Value | Units | Range |
|---|---|---|---|---|
| Parameters of the Host System, primarily sourced from [1]. | | | | |
| xS0 | extracellular substrate concentration | $10^4$ | mols | - |
| sS | stoichiometry of substrate to precursor conversion | 0.5 | - | - |
| vT | $k_{cat}$ of transport reaction | 728 | 1/(mols· min) | - |
| vE | $k_{cat}$ of host metabolic enzyme | 5800 | 1/(mols· min) | - |
| KmT | Michaelis constant of transport reaction | 1000 | mols | - |
| KmE | Michaelis constant for host metabolic enzyme | 1000 | mols | - |
| wj | max TX rate of metabolic proteins (T,H,E) | 4.14 | mols/min | - |
| wH | max TX rate of house-keeping proteins | 948.93 | mols/min | - |
| wR | max TX rate of ribosomal protein | 930 | mols/min | - |
| wr | max TX rate of ribosomal RNA | 3170 | mols/min | - |
| oj | effective Km of TX of mRNA of host enzymes (T,H,E) | 4.38 | mols | - |
| oR | effective Km of TX of ribosomal mRNA and RNA | 426.87 | mols | - |
| nj | avg no. of codons on mRNA of host enzymes (T,H,E) | 300 | no. of codons | - |
| nR | avg no. of codons on mRNA of ribosomal mRNA | 7459 | no. of codons | - |
| bj | fwd rate of active ribosomes binding with mRNA | 1 | 1/(mols· min) | - |
| uj | rate of unbinding ribosome from mRNA | 1 | 1/min | - |
| $b_\rho$ | rate of ribosomal RNA binding ribosome protein part | 1 | 1/(mols· min) | - |
| $u_\rho$ | rate of ribosomal RNA unbinding ribosome protein part | 1 | 1/min | - |
| $\delta_m$ | average mRNA degradation rate | 0.1 | 1/min | - |
| $k_H$ | inverse threshold conc of house-keeping proteins self-reg | 152219*0.8 | mols | - |
| $h_H$ | Hill coeff. of self-regulation of house-keeping proteins | 4 * 2 | - | - |
| $\gamma_{max}$ | maximal translation (TL) rate | 1260 | 1/min | - |
| $k_\gamma$ | threshold conc of precursor for half-max TL rate | 7 | mols | - |
| M0 | total cell mass | 1e8 | mols | - |
| xphi | scaling factor for max TX rate of transporter X | 0.05 | - | - |
| vX | $k_{cat}$ of native exporter X of $p_i$ | 726 | 1/(mols· min) | - |
| KmX | Km of native exporter X of $p_i$ | 1000 | mols | - |
| Parameters of TX Expression and Kinetics of Heterologous Synthesis Pathway Enzymes | | | | |
| w0 | leakiness of synth. promoters (as a frac. of their max TX) | 1e-4 | - | - |
| wEp | max TX rate of Ep | 20 | mols/min | - |
| k_Ep | $k_{cat}$ for kinetics of Ep | vE | 1/(mols· min) | - |
| Km_Ep | Km for kinetics of Ep | KmE | mols | - |
| wTp | max TX rate of Tp | 20 | mols/min | - |
| k_Tp | $k_{cat}$ for kinetics of Tp | vT | 1/(mols· min) | - |
| Km_Tp | $k_{cat}$ and Km for kinetics of Tp | KmT | mols | - |

**Supplementary Table 2:** Model parameters, table 2 of 2.

| Parameter | Description | Nominal Value | Units | Range |
|---|---|---|---|---|
| Parameters of the Inducible Genetic Control System | | | | |
| wTF | max transcription rate of TF | 20 | mols/min | - |
| sTX_T | scaling maximum TX rate of host T | 1 | - | $[10^{-3},2]$ |
| sTX_E | scaling maximum TX rate of native E | 1 | - | $[10^{-3},2]$ |
| sTX_Ep | scaling maximum TX rate of $E_p$ | 0 | - | $[10^{-3},2]$ |
| sTX_Tp | scaling maximum TX rate of Tp | 0 | - | $[10^{-3},2]$ |
| sTX_TF | scaling maximum TX rate of TF | 0 | - | $[10^{-3},2]$ |
| K_T | biosensor affinity for native T gene promoter | 1/(1000/300) | 1/mols | $[10^{-6},1]$ |
| K_E | biosensor affinity for native E gene promoter | 1/(1000/300) | 1/mols | $[10^{-6},1]$ |
| K_Ep | biosensor affinity to bind to inhibit TX of Ep | 1/(1000/300) | 1/mols | $[10^{-6},1]$ |
| K_Tp | biosensor affinity for TX of Tp | 1/(1000/300) | 1/mols | $[10^{-6},1]$ |
| K_TF | biosensor affinity for itself (TF auto-reg) | 1/(1000/300) | 1/mols | $[10^{-6},1]$ |
| $\eta$ | indicating nature of TF regulation | 0 | - | { -1,0,1} |
| Parameters of Inducer Uptake Kinetics and its Sequestering of the TF | | | | |
| kdiffI | rate of inducer diffusion in and out of a cell | 3600/60 | 1/min | - |
| VolCell | volume of cell in L (Neidhardt (1990)) | 1e-15 | L | - |
| VolCult | working volume of culture in L (3L vessel) | 1.25 | L | - |
| ksf | fwd rate of TF sequestration by inducer I | 1057.7/60 | $1/(\text{mols}^2\cdot \text{min})$ | - |
| ksr | rev. rate of TF sequestration by inducer I | 1292.1/60 | 1/min | - |
| Parameters of Batch Culture | | | | |
| $t_{max}$ | maximum of batch culture | 7*24*60 | min | - |
| $\tau$ | time of induction | - | min | [1,7*24*60] |
| $N_0$ | initial bacterial population in batch culture | $10^6$ | biomass | - |
| $S_{x,0}$ | initial glucose conc. (substrate) in media | $\frac{10*\text{VolCult}}{180}*6.02*10^{23}$ | mols | - |
| $I_0$ | concentration of added inducer | 0, else $10^{23}$ at $t=\tau$ | mols | - |

The set of parameter values in Supplementary Tables 1 and 2 are our nominal set, unless otherwise described.

#### SN2 Numerical methods for simulating the full bioprocess model

**Implementing/simulating model in MATLAB.** The ordinary differential equation model was implemented in MATLAB 2020a with dynamics simulated with the in-built stiff solver `ode15s` with tolerances of $10^{-6}$ and `NonNegative` such that no states could be negative. To improve the speed and accuracy of the solver, the Jacobian was calculated symbolically and supplied as an additionally auxillary function.

**Initial conditions for simulations of molecular dynamics within a single cell.** We defined the initial conditions for simulations of the model of the molecular dynamics within a single cell ($\dot{N(t)} = \dot{S(t)} = \dot{P(t)} = \dot{I(t)} = 0$), to steady state as: $N(t=0)=1$, $S(t=0)=10^4$, $s_i(t=0)=10^6$, $e=10^6$, $T(t=0)=E(t=0)=R(0)=100$ and $\text{TF}(t=0)=1$.

**Initial conditions for simulations of the batch culture.** We define the initial conditions for the multi-scale batch culture model as: $N(t=0) = 10^6$, $S(t=0) = 4.18 \times 10^{22}$ molecules (equivalent 10g glucose in a culture vessel of 1.25L), $P(t=0) = I(t=0) = 0$, and cellular and pathway initial conditions determined from simulations of the model of the single cell to steady state without any inducer.

**Simulating induction.** At $t = t_{\text{ind}}$, concentration of $I$ is defined as $10^{23}$ molecules, as a step input.

**Characterising performance of the controller design.** To characterise a controller design, we simulated the batch culture until all extracellular substrate is depleted ($S(t) = 0$). We define time at this point as $t_{\text{end}}$, and we calculate the yield (pY) and the volumetric productivity (vP) as:

$$\text{vP} = \frac{P(t = t_{\text{end}})}{t_{\text{end}}}, \quad \text{pY} = \frac{P(t = t_{\text{end}})}{S(t = 0)}. \tag{32}$$

#### SN3 Normalising objective values for solving the multi-objective optimisation problems

As described in the main text, our control strategy seeks to maximise yield and volumetric productivity (as defined in Equation (32) above). These objectives are often opposing, resulting in a hard-trade off. We therefore employed multiobjective optimisation routine to identify potential designs along this trade-off. In all cases models where solved numerically using MATLAB's Global Optimization Suite function `gamultiobj`. Please see main text Methods 4.3 for the definition of each of the multi-objective optimisation problems solved in this study.

The value of the objectives vary over orders of magnitude, with pY taking values between [0, 1] (by definition) and vP take values on the order of $10^{10}$-$10^{20}$. This can complicate identifying the Pareto-front as there will inherently be a biased in improving the vP values over the pY values. To ensure each objective had the same weight, we normalised each objective by its maximum value.

To find the maximum value of each objective, we applied a genetic algorithm (the `ga` function from MATLAB's Global Optimisation Toolbox (version 4.4)) to maximise vP alone and then pY alone, for a given circuit topology (defined by $g_{\text{ct}} = [g_T, g_E, g_{E_p}, g_{T_p}, g_{\text{TF}}]$, as in Equation (7) of Methods 4.3 in the main paper). We defined this optimisation problem as:

$$\left(\mathrm{vP}^{\max} = \max_{v_{\mathrm{ct}},\, \mathrm{sTX}_e,\, K_e,\, \tau} \mathrm{vP}\right) \qquad \text{or} \qquad \left(\mathrm{pY}^{\max} = \max_{v_{\mathrm{ct}},\, \mathrm{sTX}_e,\, K_e,\, \tau} \mathrm{pY}\right), \tag{33}$$

where :

$(i)$ we scale the enzyme transcription rate ($T_x(e)$) by scaling factor $\mathrm{sTX}_e$ :

$\widehat{T}_x(e) = \mathrm{sTX}_e \cdot T_x(e)$, for enzymes $(e)$ : $T$, $E$, $E_p$, $T_p$, TF,

where $10^{-3} \leq \mathrm{sTX}_e \leq 2$, where $\mathrm{sTX}_e$ denotes $\mathrm{sTX}_T$, $\mathrm{sTX}_E$, $\mathrm{sTX}_{E_p}$, $\mathrm{sTX}_{T_p}$, $\mathrm{sTX}_{\mathrm{TF}}$ ;

$(ii)$ the TF's "strength" of control on the respective enzyme, denoted $K_e$, is:

$$\begin{cases} 10^{-6} \leq K_e \leq 1 \quad \text{if} \quad g_e \neq 0\,, \\ K_e = 0 \quad \text{if} \quad g_e = 0\,, \end{cases} \qquad \text{where } K_e \text{ denotes } K_T,\, K_E,\, K_{E_p},\, K_{T_p},\, K_{\mathrm{TF}}\,;$$

$(iii)$ and the induction time $\tau$ is:

$0 \leq \tau \leq 24 * 60$ ;

where the constraints are identical to those shown in Equations (8)–(10) of the main paper.

We used these values to scale the objectives so that both vary both between [0, 1], giving each objective equal weight, each objective now re-defined as:

$$\mathrm{vP} = \frac{\mathrm{vP}}{\mathrm{vP}^{\max}} \quad \text{and} \quad \mathrm{pY} = \frac{\mathrm{pY}}{\mathrm{pY}^{\max}}. \tag{34}$$

To identify the Pareto optimal designs along any performance trade-off, between these two extremes, we applied the multi-objective genetic algorithm (the `gamultiobj` function from MATLAB's Global Optimisation Toolbox (version 4.4)) to solve the optimisation problem defined in Equations (6)–(10) of the main paper.

#### SN4 Expressing production pathway drains metabolism and reduces cell growth but increases expression of all enzymes and product synthesis

We investigated how increasing the expression of the product synthesis pathway enzyme in *a cell* ($E_p$ in Figure 17a) affects cellular processes, and how these feed back to affect product synthesis, using the model of the single host cell. We expect that increasing expression of $E_p$ will compete for ribosomes and drain host metabolite ($s_i$), and so reduce cell growth. To quantify how ribosome competition or drain of host metabolite separately impacts cell growth, we simulated cell growth with increasing transcription rates of an inactive form of the synthesis enzyme ($E_p^* = E_p(k_{\text{cat}} = 0)$) and compared to the simulated growth for increases in the transcription of an active form of the enzyme ($E_p = E_p(k_{\text{cat}} > 0)$). Results reaffirm that expressing the inactive enzyme indeed reduces cell growth at higher expression rates (Figure 17c dashed curve), due to sequestering ribosomes (Supplementary Figure 1c dashed curve). The additional draining of the host metabolite ($s_i$) from expressing the active enzyme $E_p$ reduces cell growth further (Figure 17c solid curve) because its leads to reduced transcription and translation precursor $ee$ (Supplementary Figure 1b) and a loss in total ribosomes (Figure 17d, Supplementary Figure 1c solid curve).

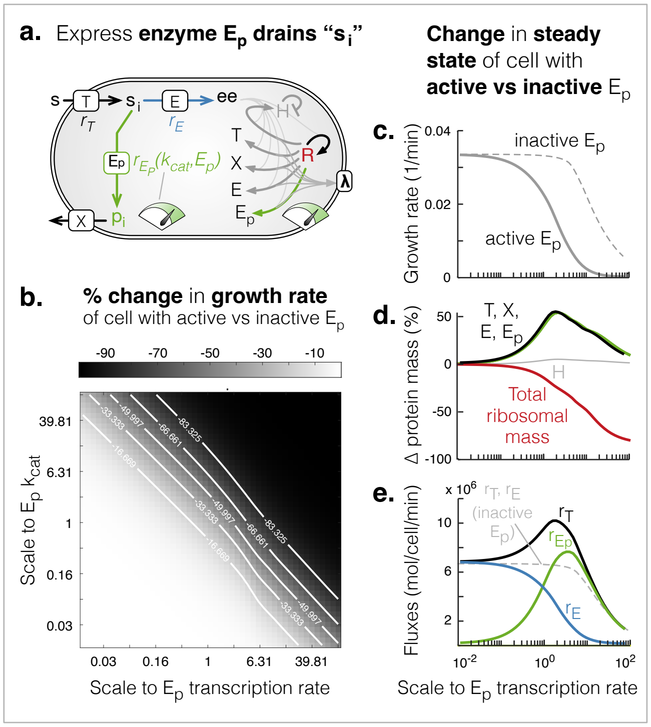


**Supplementary Figure 17: Increased draining of host metabolite can feed back to improve product synthesis.** (**a**). Schematic of the model of host and synthesis enzyme expression (gray and green arrows), host metabolism and synthesis flux (black, blue and green arrows), and growth ($\lambda$). (**b**). Heatmap of the percent 'increase' in the growth of the cell expressing the active pathway enzyme ($E_p(k_{\text{cat}} > 0)$) compared to the cell expressing the inactive enzyme ($E_p^* = E_p(k_{\text{cat}} = 0)$), for fold-changes in the maximum transcription rate and turnover rate ($k_{\text{cat}}$) of $E_p$. (**c** − **e**). Change in the steady state growth rate (**c**), percentage change in protein mass (**d**), and reaction fluxes (**e**), for fold-changes in transcription of active $E_p$ compared to inactive $E_p$.

Despite the loss in translation precursor $ee$ and ribosomes, we observed an increase in translation of all metabolic enzymes (nutrient transporter $T$, product exporter $X$, product synthesis enzyme $E_p$, host enzyme $E$) (Figure 17d). The simulations show that the loss of $ee$ causes a relatively greater loss in the transcription of ribosomal mRNA and rRNA compared to the mRNA of metabolic enzymes (Supplementary Figure 1d, e). The energy threshold (i.e. amount of $ee$ required) for ribosomal transcription ($o_R$) is higher than for enzymes ($o_X$), so as energy levels ($ee$) fall there is a larger loss in ribosomal transcription compared to the transcription of metabolic enzymes [3]. Whilst the number of ribosomes (Figure 17d) and cell growth (Figure 17c) falls, the higher proportion of mRNA of metabolic enzymes compared to ribosomal mRNA means higher fractions of ribosomes will be translating enzymes rather than ribosomes (Supplementary Figure 1e). This fundamentally shifts the cell proteome from a ribosome-dominated to an enzyme-dominated mass fraction (Figure 17d). More enzymes in turn increases the flux of nutrient uptake ($r_T$ in Figure 17e) and product synthesis ($r_{E_p}$ in Figure 17e). However, if expression of the synthesis pathway enzyme is increased to much higher rates, there is a much greater loss of ribosomes and this ends up reducing the translation of all enzymes/proteins (Figure 17d), consequently reducing nutrient uptake and product synthesis (Figure 17e).

Altogether, by activating product synthesis and inducing the "burden" of expressing the additional synthesis pathway enzyme and draining of host metabolites, the cell's natural regulation mechanisms can shift the cell into a state that favours product synthesis. However, to maximise product synthesis flux we must tune to an optimal "burden" of $E_p$ expression rate (Figure 17e), albeit at a cost to cell growth (Figure 17c).

#### SN5 Comparing the dynamics of one-sided and two-sided gene circuits

To understand how circuits with different topologies generate similar performance in terms of volumetric productivity and yield, we simulated specific designs with similar volumetric productivity ($\sim 5 \times 10^{19}$ molecules per min per culture) and compared their dynamics. For this analysis we focused on four topologies (as illustrated in Supplementary Figure 18, right): topology (A) - where the TF activates the host enzyme $E$ and inhibits the pathway enzyme $E_p$ and transporter $T_p$ (purple curves); topology (B) - where TF controls $E$ and $E_p$ in the same way as in (A) but the pathway transporter $T_p$ is consitutively expressed (blue curves); topology (C) - where the TF activates $E$ and $E_p$ is constitutively expressed (yellow curves); and topology (D) - where the TF inhibits $E_p$ and $E$ is constitutively expressed (orange curves).

Our analysis shows that topology (D) shows the greatest population growth and therefore shorter culture times while topologies (A), (B) and (C) show similar dynamics at the level of the whole culture (Supplementary Figure 18a). Topology (C) shows slower population growth due to the burden of constitutive $E_p$ production but this potential loss of volumetric productivity is countered by initial higher production of product (Supplementary Figure 18a, center and right plots).

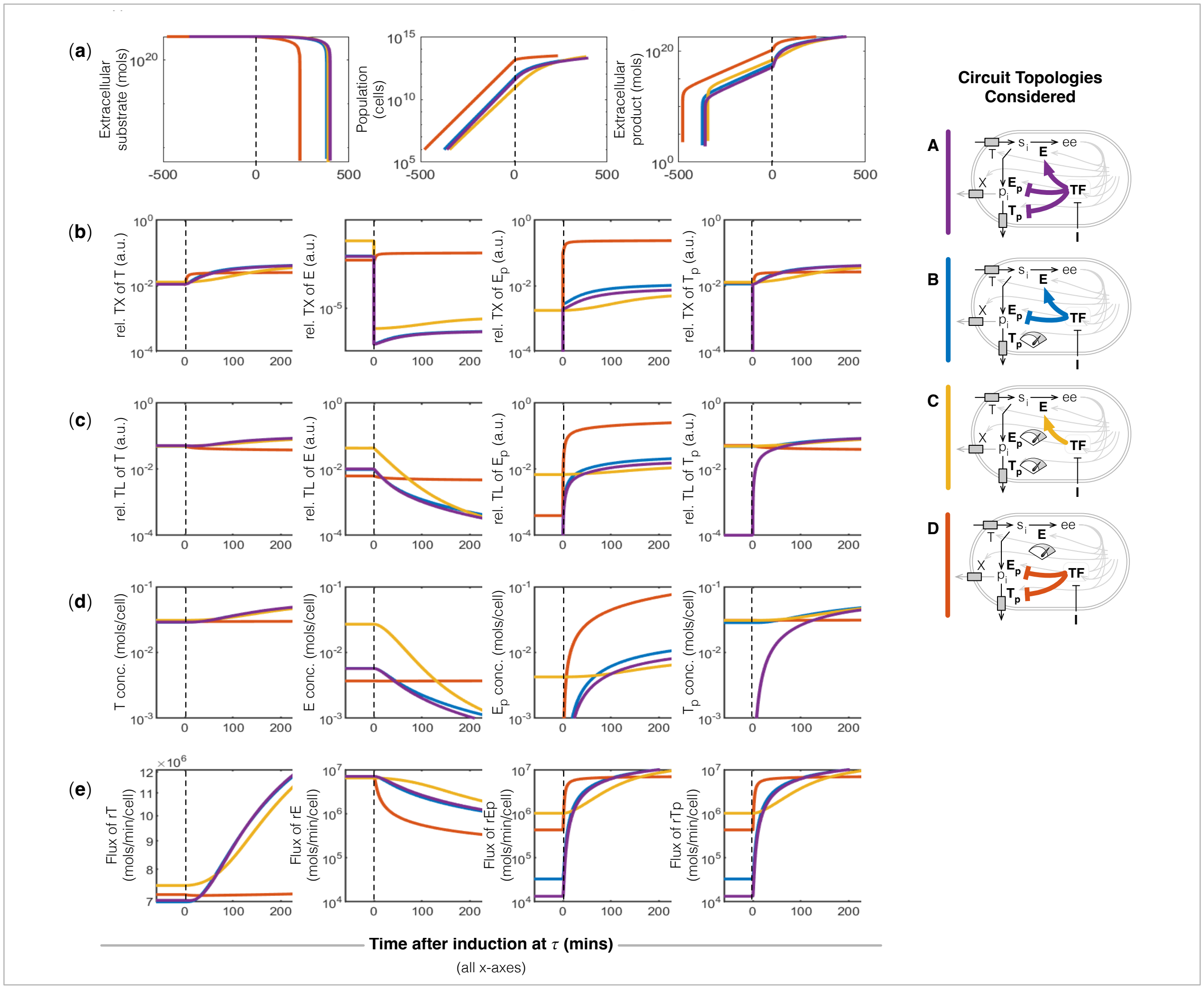

**Supplementary Figure 18: Dynamics of induced chemical production using different circuit topologies (illustrated on the right) with similar performance.** Plots of the dynamic response after adding inducer at time point 0, of culture level variables, including external substrate utilization (**a**, first), population (**a**, second) and external product accumulation (**a**, third); and the transcription (TX) rates relative to total transcription (**b**), translation (TL) rates relative to total translation (**c**), and concentrations (**d**) of enzymes $T$, $E$, $E_p$, $T_p$. (**e**). Flux of reactions catalysed by enzymes $T$, $E$, $E_p$, $T_p$.

Upon induction at time $\tau$, the relative transcription rate of $E$ falls for topologies (A), (B) and (C) due to action of the controller. However, for topology (D), we see an unintended increase in the relative transcription rate of the host enzyme $E$ (Supplementary Figure 18b). As the cell's metabolism is diverted to the pathway, the amount of energy $ee$ falls and due to the natural regulation in the model, the transcription rates of enzymes in the model are increased (see detailed analysis of this phenomenon in Supplementary Note SN4). Therefore, by not directly inactivating or inhibiting $E$'s transcription, the indirect regulatory feedback from resource competition increases the transcription of $E$ (Supplementary Figure 18b, second plot). While $E_p$ is not directly regulated in topology (C), the same proteome reallocation leads to an increase in its transcription after the induction switch is activated (Supplementary Figure 18b, third plot). The same indirect activation causes an increase in $T_p$ production in topology (B) without direct regulation (Supplementary Figure 18b, fourth plot). In all topologies, the unregulated host transporter $T$ rises (Supplementary Figure 18b, first plot).

The relative translation rates (i.e. the translation rate as a proportion of total translation) shows similar dynamics (Supplementary Figure 18c). Both topology (A) and (B) show similar translation dynamics across host ($T$, $E$) and pathway ($E_p$, $T_p$) genes. The increase in mRNA of host enzyme $E$ in topology (D) means that translation of $E$ is maintained (Supplementary Figure 18c), therefore for optimal production higher amounts of $E_p$ is required (Supplementary Figure 18d). This results in reduced translation of $T$ and $T_p$ which reduces the systems input and output fluxes and reduces system productivity (Supplementary Figure 18c). Despite $E_p$ being unregulated in topology (C), the increase in transcription results in an slight increase in translation upon system activation (Supplementary Figure 18c). In all topologies except (D), the unregulated host transporter $T$ and synthetic exporter $T_p$ translation rate rises (Supplementary Figure 18c).

These changes in transcription and translation are reflected as expected in the concentrations of the key host and pathway enzymes (Supplementary Figure 18d). Topologies (A) and (B) show similar dynamics for all proteins. Whilst $T_p$ is initial present in topology (B), where it is unregulated the non-regulatory interactions described above result in an increase in $T_p$ after the switch is activated to a similar mass fraction as in Topology (A), where the gene directly regulated (Supplementary Figure 18d, fourth plot). The cause of the slightly lower volumetric productivity of topology (C) compared to (A) and (B) is driven by the lower $E_p$ (Supplementary Figure 18d, third plot). Both host enzyme $E$ and pathway enzyme $E_p$ are more highly expressed which reduces growth rate and population size before induction (Supplementary Figure 18a, second plot). The lack of reduction of host enzyme $E$ in topology (D), means to switch the cells to production with high productivity requires high expression of $E_p$ (in fact, mass fraction of $E_p$ in this design is over an order of magnitude higher that of the other designs, Supplementary Figure 18d, third plot). This prevents burden induced natural proteome reallocation from increasing host $T$ or pathway $T_p$ (Supplementary Figure 18d, first and fourth plots).

These protein's changes are reflected in pathway fluxes (Supplementary Figure 18e): upon activation of the switches, in all topologies bar (D), show an increase in substrate import through $T$ and product export through $T_p$ (Supplementary Figure 18e, first and fourth plots). The higher production of $E_p$ for topology (D) prevents this as described above. Over all, we see that topologies (A) and (B) have similar flux dynamics (Supplementary Figure 18e). The fluxes of (C) show qualitatively similar dynamics to (A) despite the much more simple regulation in the circuit (Supplementary Figure 18e, yellow compared to purple curves).
